# Direct genome editing of patient-derived xenografts using CRISPR-Cas9 enables rapid *in vivo* functional genomics

**DOI:** 10.1101/588921

**Authors:** Christopher H. Hulton, Emily A. Costa, Nisargbhai S. Shah, Alvaro Quintanal-Villalonga, Glenn Heller, Elisa de Stanchina, Charles M. Rudin, John T Poirier

## Abstract

Patient-derived xenografts (PDXs) constitute a powerful set of preclinical models for *in vivo* cancer research, reflecting the spectrum of genomic alterations and therapeutic liabilities of human cancers^1-4^. In contrast to either cancer cell lines or genetically engineered mouse models, the utility of PDXs has been limited by the inability to perform targeted genome editing of these tumors. To address this limitation, we have generated a lentiviral platform for CRISPR-Cas9 editing of PDXs using a tightly regulated, inducible Cas9 vector that does not require *in vitro* culture for selection of transduced cells. We demonstrate the utility of this platform in PDXs (1) to analyze genetic dependencies by targeted gene disruption and (2) to analyze mechanisms of acquired drug resistance by site-specific gene editing using templated homology-directed repair. This flexible system has broad application to other explant models and substantially augments the utility of PDXs as genetically programmable models of human cancer.

---

PDXs recapitulate the complex genotypes and intratumoral heterogeneity of their tumors of origin and are not subject to the selective pressures imposed by *in vitro* cell culture since they are maintained exclusively *in vivo*^5-7^. These features have driven the rapid adoption and widespread use of PDXs in preclinical and co-clinical drug development, evaluation of biomarkers and imaging agents, and mechanistic investigation of acquired treatment resistance^8-10^. PDXs have also proven to be valuable models of tumor types or variants for which *in vitro* models are not readily available^11,12^.

The ability to genetically manipulate cancer models has played an essential role in defining the functional contributions of individual genes and variants. CRISPR-Cas9 genome editing has greatly expanded our ability to rapidly analyze how specific genes and genotypes contribute to carcinogenesis and tumor maintenance^13,14^. CRISPR-Cas9 can be used to disrupt genes through the introduction of frameshift insertions and deletions (indels) by non-homologous end joining or to make precise genomic alterations through homology-directed repair (HDR)^15^. However, *in vivo* applications of CRISPR-Cas9 genome editing have been restricted to xenografts of established human cell lines and genetically engineered mouse models (GEMMs) due largely to the technical challenges imposed by the continuous *in vivo* passaging of PDXs and the need for alternative selection methods. Given the increasing role of PDXs in cancer research, a technological advance is needed to enable direct genome editing of PDXs using CRISPR-Cas9 to further our understanding of cancer biology and facilitate the development of new therapeutic strategies.

Tight temporal control of Cas9 activity in the cells that make up established tumors is essential to validate genes required for tumor maintenance and to credential suppressor mutations that may play a role in acquired drug resistance. Several inducible systems have been developed to regulate Cas9 activity at the post-translational level, yet these systems invariably suffer from aberrant or reduced Cas9 activity [reviewed in ^16^]. Doxycycline (dox)-inducible expression of Cas9 provides a combination of maximum cutting efficiency in the “on” state while minimizing Cas9 activity in the “off” state through tight transcriptional regulation. However, current systems lack complete transcriptional control by dox and are not amenable to use in PDXs because they either rely on inefficient knock-in approaches^17-19^ or employ vectors that exceed the lentiviral packaging limit and consequently result in low viral titers and predictably poor transduction efficiency^20,21^. Importantly, all current modalities of inducible Cas9 expression lack a suitable selectable marker for use in PDXs.

To overcome these limitations and enable *in vivo* functional genomic studies in PDXs, we developed pSpCTRE, an all-in-one dox-inducible Cas9 lentiviral vector tailored for use in PDXs (Fig. 1a). Leaky Cas9 expression is reduced by the placement of a TRE^3GS^ promoter in the reverse orientation relative to the viral LTRs^22^ and through use of the rtTA-V10 variant^23^. Since lentiviral titer decreases as a function of proviral size^24^, several approaches were taken to minimize the size of pSpCTRE. A minimal EFS promoter is used to drive constitutive expression of rtTA-V10 as well as a truncated human CD4 selectable marker (hereafter CD4^T^), with the latter enabling rapid, positive selection of pSpCTRE-transduced cells without intervening *in vitro* culture. These optimizations collectively result in a final SpCTRE size of 9,023 bp between the LTRs.

**Figure 1.**
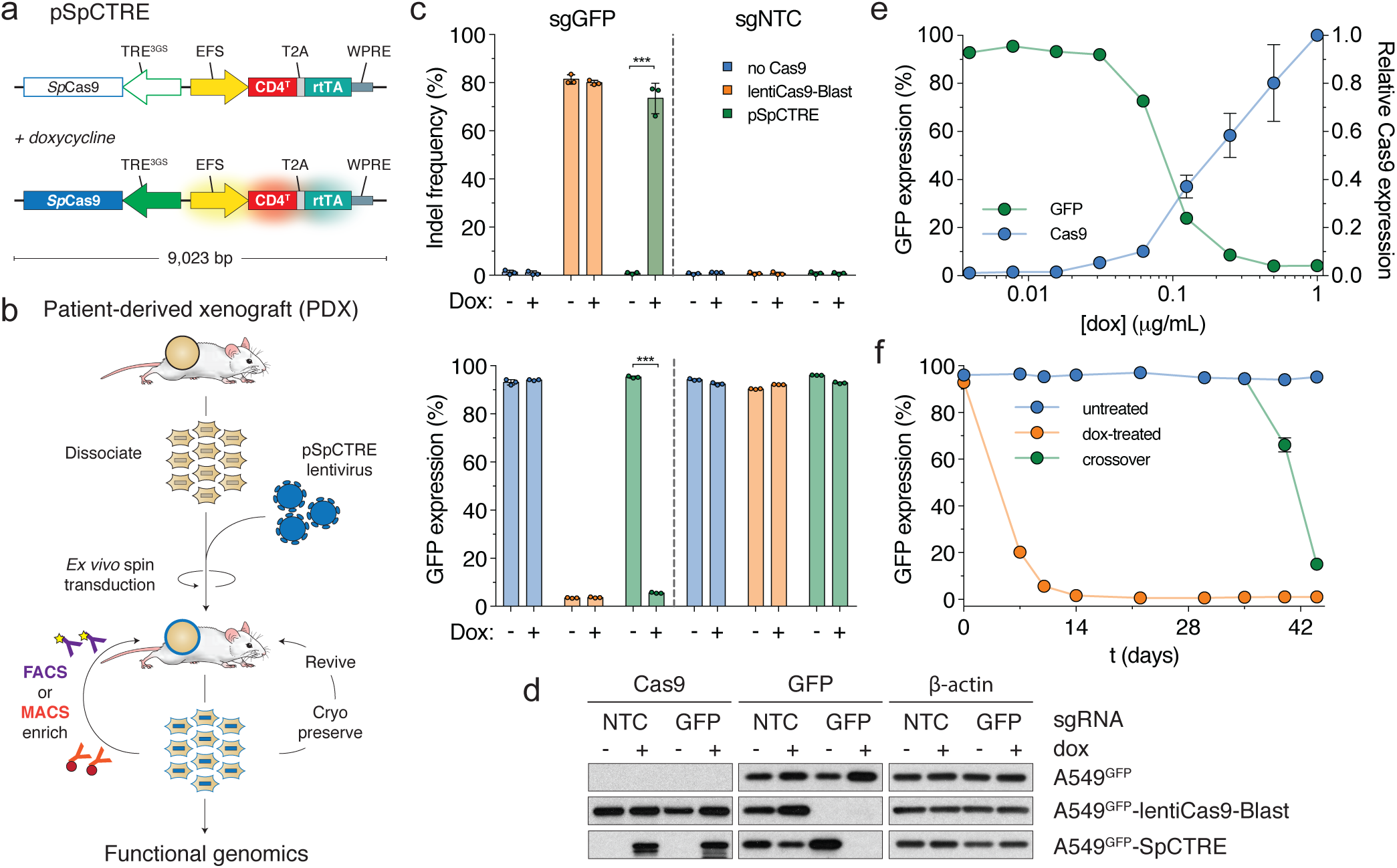
Development and validation of pSpCTRE, a lentiviral Cas9 vector for doxycycline (dox)-inducible genome editing in PDXs. (**a**) Vector map. Promoter activity and gene expression in the absence (top) or presence (bottom) of dox is depicted as none (outline), active (solid), and induced (glow). CD4^T^ is a minimal, truncated CD4 selectable marker linked to rtTA-V10 by a T2A ribosome skipping sequence (see **Supplementary Fig. 1** for CD4^T^ details). With dox, the TRE^3GS^ promoter is activated while secondarily inducing EFS promoter activity (see **Supplementary Fig. 2** for additional data). (**b**) Schematic of Cas9 PDX generation. Dissociated tumors are transduced *ex vivo* with pSpCTRE lentivirus and immediately engrafted in mice without intervening *in vitro* culture. Resultant SpCTRE PDXs (blue outline) are then subjected to CD4 enrichment, cryopreservation, or functional studies. (**c**) *In vitro* genome editing of A549^GFP^ cells lacking Cas9 (blue), with constitutive Cas9 expression from lentiCas9-Blast (orange), or with inducible Cas9 expression from SpCTRE (green) and the indicated sgRNAs. Cells were analyzed for indel frequency (top) and GFP expression (bottom). Error bars are SD (n = 3), SD < plotting character not drawn. *** p < 0.001 between groups by two-way ANOVA. NTC, Nontargeting control (**d**) Representative Western blot from panel **c**. (**e**) Dox dose response of Cas9 expression and GFP editing in A549^GFP^-SpCTRE cells with sgGFP. Error bars are SD (n = 3), SD < plotting character not drawn. (**f**) Long-term culture of A549^GFP^-SpCTRE cells with sgGFP untreated (blue) or dox-treated (orange). At day 35, previously untreated cells were crossed over to dox media (green). Error bars are SD (n = 3), SD < plotting character not drawn.

The optimal composition of CD4^T^ was empirically determined through a deletion series of the extracellular Ig-like domains required for binding of commercial α-CD4 antibodies (Supplementary Fig. 1a). While several α-CD4 antibody epitopes reside within domain D_1_^25^, we found that both domains D_1_ and D_2_ were required for binding of these antibodies (Supplementary Fig. 1a). Given this finding, CD4^T^ is composed of the CD4 signal peptide and extracellular domains D_1_ and D_2_ fused to the transmembrane domain (TM) via a flexible linker, with the intracellular portion of the protein omitted entirely (Supplementary Fig. 1a). The final coding sequence is approximately half the size of full-length CD4 and can be used with commercial flow cytometry or magnetic bead sorting reagents (Supplementary Fig. 1b, c). Notably, in CD4-expressing cells, the absence of extracellular domains D_3_ and D_4_ in this construct allows for differentiation of CD4^T^ from full-length CD4 by counterstaining with a domain D_3_ specific antibody (Supplementary Fig. 1d).

In addition to mitigating leaky Cas9 expression, reverse orientation of the TRE^3GS^ promoter places it proximal to the constitutive EFS promoter. This promoter topology is unidirectional in the absence of dox and potently induced bidirectionally in the presence of dox, markedly increasing CD4^T^ expressed from the EFS promoter (Fig. 1a and Supplementary Fig. 2a). A feed forward loop is created by a concurrent increase in rtTA-V10 expression, resulting in an approximately 100-fold induction of CD4^T^ staining intensity over basal levels and three distinct populations of cells: 1) CD4^T^ negative, 2) CD4^T^ positive, and 3) CD4^T^ induced (Supplementary Fig. 2b, c). CD4^T^ induced expression is a robust marker of cells that express Cas9 and are competent to undergo Cas9-mediated genome editing (Supplementary Fig. 2d). This feature of pSpCTRE provides a novel cell surface marker of dox-induced transgene expression and allows for identification of cells with efficient and heritable induction of Cas9, enabling the study of heterogeneous tumor cell populations by eliminating the need for selection of single cell clones.

Cas9-expressing SpCTRE PDXs can be generated from pSpCTRE lentivirus in as few as two *in vivo* passages (Fig. 1b). In the first passage, dissociated tumors are subjected to an *ex vivo* spin transduction and immediately re-engrafted for expansion *in vivo*. Successfully transduced cells are enriched from the ensuing tumor with α-CD4 positive selection and either cryopreserved or passaged for subsequent use in functional genomic studies.

We initially examined SpCTRE editing efficiency through *in vitro* disruption of GFP in A549 cells with a single copy of GFP (A549^GFP^). In the absence of dox, A549^GFP^-SpCTRE cells transduced with sgGFP maintained GFP expression and had no detectable Cas9 protein by Western blot (Fig. 1c, d). Conversely, after exposure to dox, A549^GFP^-SpCTRE cells efficiently edited GFP, similar to A549^GFP^ cells constitutively expressing Cas9 from the lentiCas9-Blast vector (Fig. 1c, d). We observed a dose-dependent increase in Cas9 expression and CD4^T^ induction with concomitant loss of GFP expression with as little as 0.0625 µg/mL dox and a maximum loss of GFP expression at 0.25 µg/mL dox, defining an upper and lower bound for dox response (Fig. 1e and Supplementary Fig. 2b). To more sensitively screen for leaky expression of Cas9 and undesired genome editing, we cultured A549^GFP^-SpCTRE cells transduced with sgGFP in the absence of dox. We observed no decrease in GFP-positive cells over >10 passages, yet these cells retained the capacity to efficiently disrupt GFP upon dox induction after 35 days of continuous culture (Fig. 1f). These data indicate that pSpCTRE is a tightly regulated dox-inducible Cas9 vector, suggesting that this system is well suited for *in vivo* use in heterogeneous PDX tumors.

We next sought to generate a library of SpCTRE PDXs by transducing 20 PDXs representing multiple lung cancer subtypes with pSpCTRE lentivirus. We were able to detect and enrich CD4^T^ positive cells in six of these models (Supplementary Fig. 3a). The median time to establish these SpCTRE PDXs following transduction was 154 days (with a range of 108 to 305 days) and one to three *in vivo* passages were required (Supplementary Fig. 3b). Importantly, these SpCTRE PDX tumors express Cas9 exclusively when the mice are administered dox (Supplementary Fig. 3c).

We first interrogated genetic dependencies of SpCTRE PDXs using a competition assay, which allows for measurement of gene disruption effects without prior selection for cells transduced with sgRNAs (Fig. 2a). To deliver each sgRNA, we generated a fluorescent reporter lentiviral vector (sgTrack) that constitutively expresses either mCherry or GFP along with an sgRNA and can be produced at high titer. We independently transduced SpCTRE PDXs *ex vivo* with sgTrack vectors encoding a non-targeting sgRNA or an sgRNA targeting a gene of interest and mixed both populations before engrafting in mice. The fluorescent reporters associated with each sgRNA allowed us to measure the relative abundance of sgRNAs in a population as a proxy for the relative fitness of cells engineered to disrupt a gene of interest. This approach robustly identified differential fitness of cells *in vitro*, reflected in the reduction of cells carrying an sgRNA targeting the essential gene *RPA1* or the *KRAS* proto-oncogene in a KRAS^G12S^-mutant lung adenocarcinoma cell line (Supplementary Fig. 4a-c).

**Figure 2.**
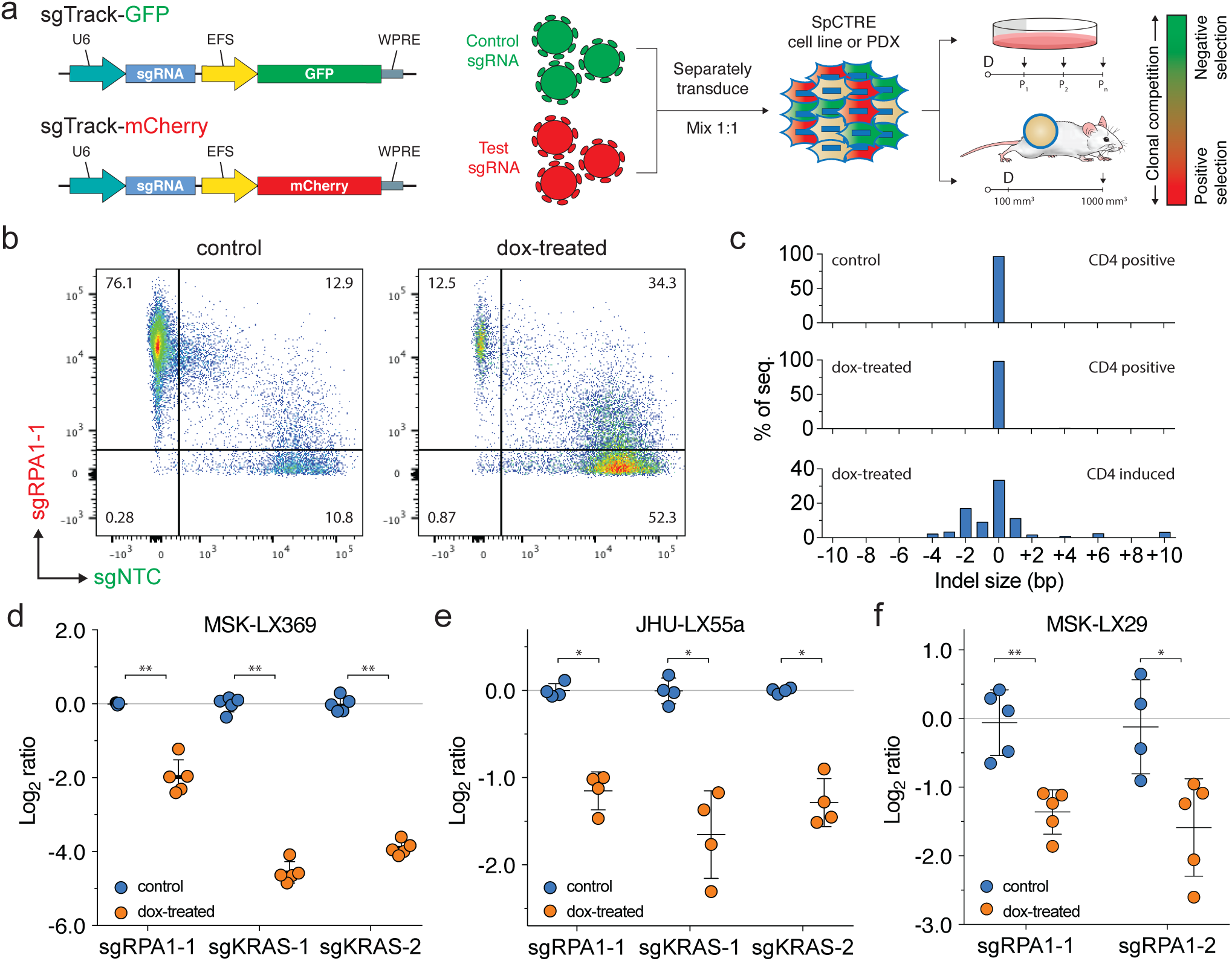
Interrogation of genetic dependencies in SpCTRE PDXs using a competition assay. (**a**) Schematic of competition assay design. Control or test sgRNAs in sgTrack fluorescent reporter lentiviral vectors are transduced independently into SpCTRE PDXs or cell lines and then admixed and either maintained in culture (if cell lines) or engrafted subcutaneously (if PDXs). Dox addition (D) and flow cytometry analysis (arrows) occurs at the indicated time points and the relative abundance of GFP and mCherry single positive cells is compared between control and dox-treated samples. (**b**) Flow cytometry analysis of representative tumors from a competition assay with sgRPA1-1 (mCherry) and sgNTC (GFP) in the MSK-LX369 lung adenocarcinoma PDX. Control tumors were gated on CD4^T^ positive cells and dox-treated tumors were gated on CD4^T^ induced cells (see **Supplementary Fig. 5** for gating strategy). (**c**) RPA1 genome editing of representative tumors from panel **b**. Analysis was restricted to sgRPA1-1 containing cells from the indicated treatment groups and CD4^T^ gates. (**d-f**) Competition assays with the indicated sgRNAs in the SpCTRE PDXs (**d**) MSK-LX369, (**e**) JHU-LX55a, and (**f**) MSK-LX29. The log ratio compares the fitness score of dox-treated mice to the fitness score of control mice (see **Supplementary Fig. 4b** for details). Error bars are SD (n = 5), SD < plotting character not drawn. Each data point represents one tumor. * p < 0.05 and ** p < 0.01 between groups by Wilcoxon rank sum test.

We next performed similar competition assays *in vivo* to study the effects of single gene disruption on the fitness of three SpCTRE PDXs. Targeting RPA1 in MSK-LX369, JHU-LX55a, and MSK-LX29, we consistently observed a depletion of cells carrying the RPA1 sgRNA and an enrichment of cells carrying a non-targeting sgRNA in CD4^T^ induced cells from dox-treated mice (Fig. 2b, d-f and Supplementary Fig. 5a-c). We verified targeted editing of the RPA1 locus in isolated tumor cells and confirmed the presence of indels exclusively in sgRPA-transduced cells with CD4^T^ induced expression (Fig. 2c). Additionally, we investigated the KRAS dependency^26,27^ of two KRAS-mutant lung adenocarcinoma SpCTRE PDXs, MSK-LX369 and JHU-LX55a. We observed a significant depletion of cells harboring KRAS sgRNAs in CD4^T^ induced cells from dox-treated mice, indicating that these PDXs are indeed KRAS dependent (Fig. 2d, e and Supplementary Fig. 5c, d). These experiments verify that this system is suited to functionally interrogate PDX gene essentiality *in vivo*.

To interrogate a broader spectrum of single nucleotide variants and compound genetic alterations found in human cancer, we generated a recombinant adeno-associated virus (rAAV) vector that delivers an sgRNA and a homology-directed repair template encoding multiple desired genetic alterations (Fig 3a). To validate this approach in SpCTRE PDXs, we chose to explore mechanisms of acquired EGFR inhibitor resistance in lung adenocarcinomas. MSK-LX29 was derived from a patient with a homozygous EGFR^L858R^ mutation who developed resistance to the 1^st^ generation EGFR inhibitor erlotinib through acquisition of a focal *MET* amplification. This PDX is resistant to single agent treatment with the 3^rd^ generation EGFR inhibitor osimertinib as well as the MET inhibitor crizotinib but is profoundly and durably sensitive to a combination therapy of the two drugs (Supplementary Fig. 6a, b).

**Figure 3.**
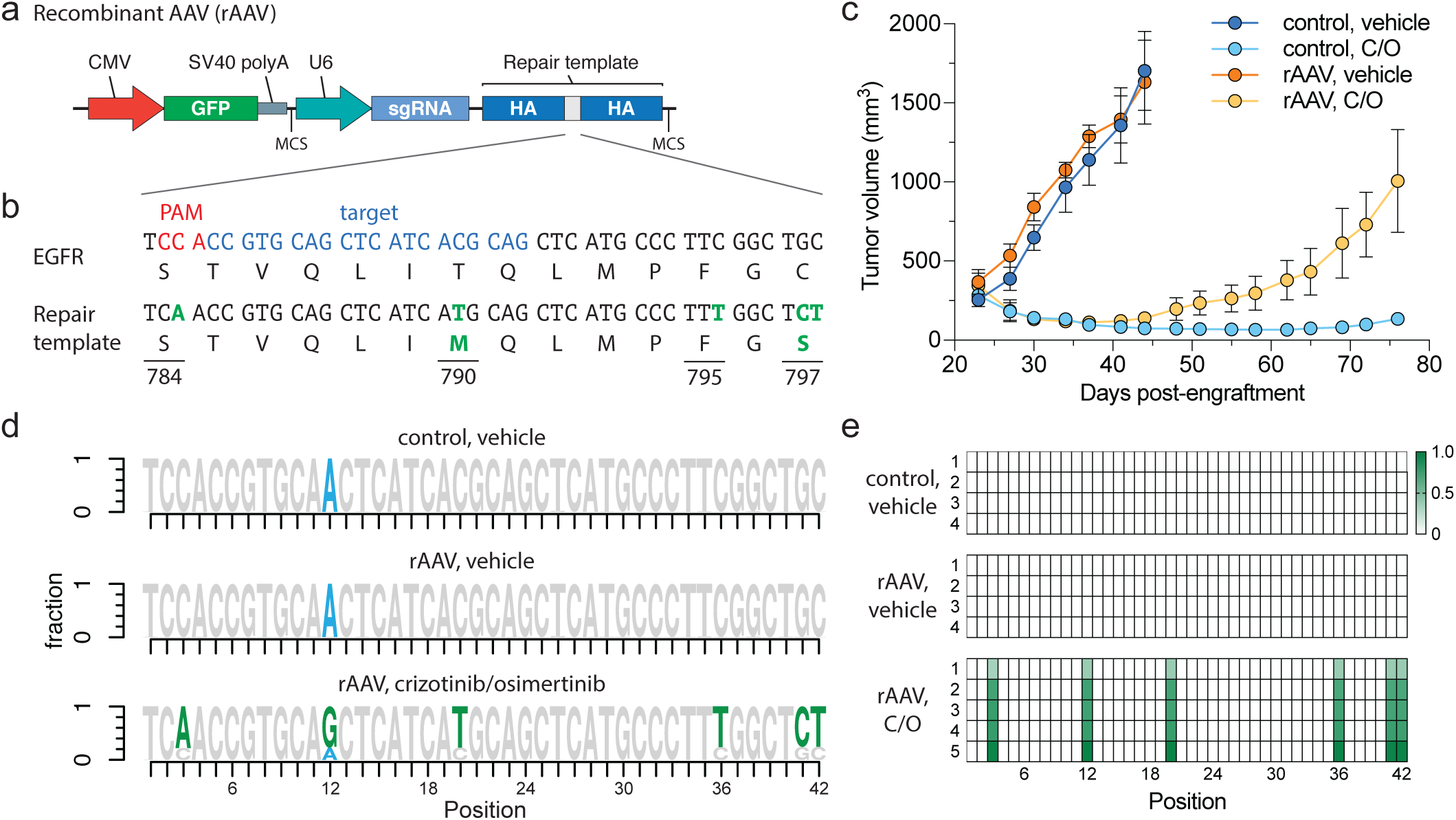
Introduction of complex drug resistance mutations in SpCTRE PDXs using recombinant AAV (rAAV). (**a**) Schematic of rAAV vector that delivers a GFP marker, an sgRNA expressed from a U6 promoter, and a homology-directed repair template that encodes mutations of interest surrounded by homology arms (HA). The U6-sgRNA and repair template are flanked by multiple cloning sites (MCS). (**b**) sgRNA target site and EGFR T790M/C797S repair template to introduce the indicated base changes (green). (**c**) Tumor volumes of MSK-LX29-SpCTRE with or without rAAV and treated with a crizotinib/osimertinib (C/O) combination or vehicle. Error bars are SD (n = 5), SD < plotting character not drawn. (**d**) Sequencing analysis of representative tumors from the indicated treatment groups. Mutations introduced by rAAV are highlighted (green). MSK-LX29 contains a homozygous SNP (blue) that is removed by the rAAV. (**e**) Heatmap depicting frequency of rAAV introduced mutations for tumors from the indicated treatment groups.

We generated an rAAV vector that encodes both an sgRNA targeting a sequence proximal to the hydrophobic binding pocket of the EGFR kinase domain and a 1.5-kb repair template centered on this region. The repair template encodes base changes that silently destroy the sgEGFR PAM sequence to prevent Cas9 re-cutting after repair and introduces *in cis* 1) a T790M 1^st^ generation EGFR inhibitor gatekeeper mutation, 2) a C797S mutation that abolishes activity of 3^rd^ generation covalent EGFR inhibitors and 3) a silent landmark mutation to mark desired base changes that originate from the repair template (Fig. 3b). Efficient generation of osimertinib resistance in EGFR-mutant PC9 cells using this template was confirmed *in vitro* (Supplementary Fig. 7a-c). While initially sensitive to a crizotinib-osimertinib combination therapy, within 2 weeks after the start of treatment MSK-LX29-SpCTRE infected with this rAAV developed resistance in all five tumors in this treatment group (Fig. 3c and Supplementary Fig. 6c). All five resistant tumors demonstrated efficient editing and selection for the templated C797S mutation (Fig. 3d, e). We conclude that an acquired resistance mutation in EGFR alone is sufficient to restore resistance to crizotinib-osimertinib combination therapy in EGFR mutant lung adenocarcinomas that acquire bypass resistance through *MET* amplification. More generally, these data confirm that pSpCTRE can be used to introduce template-directed site-specific mutations *in vivo*.

Together, this suite of vectors constitutes a core enabling technology platform for *in vivo* functional genomics in PDXs, allowing interrogation of gene essentiality, candidate drug targets, mechanisms of acquired resistance, tumor suppressor function, chemical:genetic interactions, and variants of unknown significance in this important tumor model.

## METHODS

### Patient-derived xenografts (PDXs)

All animal experiments were approved by the Memorial Sloan Kettering Cancer Center (MSKCC) Animal Care and Use Committee. Primary tumors and whole blood samples collected for generation of PDX models were obtained with informed consent from patients under protocols approved by the MSKCC and Johns Hopkins institutional review boards. Subcutaneous flank tumors were generated as described previously^9^.

#### Cloning and plasmids

The plasmids generated in this study are available on Addgene as indicated. A list of all primers used in this study is available in **Supplementary Table S1**.

pSpCTRE (Addgene plasmid # 114010): EFS promoter expresses CD4^T^-2A-rtTA-V10; TRE-3GS promoter controls tetracycline-inducible expression of *S*.*pyogenes* Cas9 as an all-in-one Tet-On system. This vector was derived from the pLVX-TetOne-Puro vector (Clontech), which was digested with XhoI and KpnI to remove the existing rtTA and Puro selection cassettes. The EFS-CD4^T^-2A-rtTA-WPRE cassette was generated by gene synthesis (IDT) and inserted into the digested vector by Gibson assembly (NEB). Cas9 was PCR amplified and inserted into the multiple cloning site by digestion with EcoRI and BamHI followed by overnight ligation with T4 ligase (NEB).

sgTrack-Gateway (Addgene plasmid # 114011), sgTrack-GFP (Addgene plasmid # 114012) and sgTrack-mCherry (Addgene plasmid # 114013): U6 promoter expresses a single sgRNA; EFS promoter upstream of the Gateway cassette expresses either TurboGFP or mCherry in the respective reporter vectors. These vectors are derived from lentiCRISPRv2^28^, wherein Cas9 was replaced by a Gateway cassette in-frame with the existing 2A-Puro by PCR amplification of the Gateway cassette and Gibson assembly into an XbaI and BamHI digested backbone. Gateway donor vectors with closed TurboGFP and mCherry cDNAs were used to generate the respective reporter vectors using LR clonase (Invitrogen) according to the manufacturer’s protocol. sgRNAs were cloned into these vectors as previously described^28^.

pAAV-GFP-sgHDR: pAAV-GFP was a gift from John T Gray (Addgene plasmid # 32395)^29^. A synthetic DNA fragment comprising an SV40 polyA terminator, partial multiple cloning site (BglII, HindIII), the human U6 promoter, EGFR sgRNA, a 1.5 kb EGFR homology directed repair template containing the T790M mutation and destroying the sgEGFR PAM sequence, and a second partial multiple cloning site (ClaI, XhoI, XbaI) was cloned into the NotI and XbaI restriction enzyme sites of pAAV-GFP, replacing the original beta-globin polyA fragment. EGFR C797S coding and silent repair landmark mutations were inserted via site directed mutagenesis using primer pair P1 to generate the rAAV vector.

lentiCas9-Blast (Addgene plasmid # 52962) and lentiGuide-Puro (Addgene plasmid # 52963) were gifts from Feng Zhang. pMD2.G (Addgene plasmid # 12259) and psPAX2 (Addgene plasmid # 12260) were gifts from Didier Trono. pLenti CMV Neo DEST (705-1) (Addgene plasmid # 17392) and pLenti CMV Puro (w118-1) (Addgene plasmid # 17452) were gifts from Eric Campeau and Paul Kaufman^30^. All Gateway recombination reactions were performed with BP clonase or LR clonase used according to the manufacturer’s protocol (Invitrogen). TurboGFP cDNA was cloned into pLenti CMV Neo DEST (705-1). pDONR221/CD4 was purchased from DNASU (HsCD00413471). CD4ΔD_2-4_;ΔIC linked to rtTA-V10 by a T2A sequence was generated by gene synthesis (IDT), Gateway adapted by PCR using primer pair P2, and cloned into pDONR221. CD4 domain D_2_ was then added to this construct to generate CD4ΔD_3-4_;ΔIC (CD4^T^) by performing an outward PCR with primer pair P3, extracting domain D_2_ from pDONR221/CD4 using primer pair P4 and performing a Gibson assembly (NEB) with the PCR products. All CD4 constructs were then cloned into pLenti CMV Puro (w118-1). sgRNAs were cloned into lentiGuide-Puro as previously described^28^. All transformations were performed in One Shot Stbl3 Chemically Competent cells (Invitrogen). Plasmids were purified using QIAquick Spin Miniprep or Plasmid Plus Midi kits (Qiagen) and digest verified prior to use. All PCRs were performed with Phusion High-Fidelity PCR Master Mix with HF Buffer (NEB).

#### sgRNA sequences

Target sequences for sgRNAs used in this study are available in **Supplementary Table S2**.

#### Cell culture and lentivirus production

A549, PC9, and HEK239T cells were purchased from ATCC. A549 and PC9 cells were maintained in RPMI-1640 media supplemented with 10% Tet-Free FBS (Gemini) and 1x penicillin/streptomycin (pen/strep, Gibco) and HEK239T cells were maintained in DMEM media supplemented with 10% FBS (Gemini) and 1x pen/strep. All cell lines were verified negative for mycoplasma within 6 months of use. Lentivirus was produced by transfecting HEK293T cells with a 3:2:1 ratio of lentiviral plasmid:psPAX2:pMD2.G with JetPrime transfection reagent (Polyplus) at a 2:1 JetPrime:DNA ratio. Media was changed 24 h after transfection and viral supernatants were collected 72 h after transfection. Viral supernatants were syringe filtered with a 0.45 uM PVDF filter (Millipore) and concentrated approximately 20 fold with Lenti-X Concentrator (Clontech) according to the manufacturer’s protocol. All lentivirus was titered in A549 cells to control for batch-to-batch variability and to normalize titers between different lentiviral backbones. *In vitro* lentiviral transductions were performed with 8 µg/mL hexadimethrine bromide (polybrene, Sigma) and at a multiplicity of infection (MOI) of approximately 1, unless otherwise stated.

#### *In vitro* validation of pSpCTRE

An A549^GFP^ cell line containing a stably integrated, single-copy TurboGFP gene was generated by transducing A549 with pLenti CMV Neo/TurboGFP lentivirus (MOI 0.3) and single cell FACS sorting using a FACSAria (BD Biosciences). A549^GFP^ cells were subsequently transduced with lentiCas9-Blast or pSpCTRE lentivirus (MOI 0.3) and selected with blasticidin or CD4^T^ single cell FACS sorting, respectively, to generate stable A549^GFP^ Cas9 cell lines. A549^GFP^ Cas9 cells were transduced with lentiGuide-Puro/sgGFP or sgNTC lentivirus and selected with puromycin for 3 days. For analysis of GFP editing efficiency and dox dose response of pSpCTRE, 1 x 10^5^ cells were plated in a 10 cm plate and treated with 0.5 µg/mL doxycycline (dox, Sigma) or a dox dose range, respectively, for 10 days. Cell pellets were collected to perform Western blot, Tracking of Indels by Decomposition (TIDE), or flow cytometry analysis as described below. To screen for Cas9 editing in the absence of dox, A549^GFP^-SpCTRE cells with lentiGuide-Puro/sgGFP were maintained in culture for 6 weeks in RPMI with tet-free FBS. Cells were split when they reached approximately 70% confluence and flow cytometry analysis was performed every 7 days as described below. At day 35, cells not previously treated with dox were split and 0.5 µg/mL dox was added to half of the cells (crossover). *In vitro* competition assays were performed in A549 cells transduced with lentiCas9-Blast or pSpCTRE lentivirus (MOI 0.3) and selected with blasticidin or CD4^T^ single cell FACS sorting, respectively. A549 Cas9 cell lines were independently transduced with sgTrack lentivirus and, after 3 days, cells containing control and test sgRNAs were mixed with an equal ratio of GFP and mCherry positive cells. Cells were split and flow analysis was performed as described below every 4 days. Results for all experiments represent three independent biological replicates.

#### Antibodies

A list of antibodies used in this study is available in **Supplementary Table S3**.

### Protein extraction, **Western blotting and LiCor protein quantification**

Whole cell lysates were prepared from frozen cell pellets or flash frozen tumor samples using RIPA lysis buffer with 1x HALT protease inhibitor cocktail (Thermo). Cell pellets were resuspended in 5 volumes of cold lysis buffer and incubated on ice for 10 minutes, followed by sonication for 10 seconds with a 200V microtip sonicator set to 40% amplitude (QSonica, CL 18). Lysates were clarified by centrifugation at 20,000 x g for 10 minutes at 4C. Protein extraction from flash frozen tumor samples was performed as previously described^9^. Protein was quantified using a BCA protein assay kit (Pierce) and samples were denatured at 70C for 10 minutes in NuPAGE LDS sample buffer with NuPAGE sample reducing agent and then resolved on a 4-12% Bis-Tris gradient gel (Invitrogen). For chemiluminescent detection, gels were wet-transferred to 0.45 µm Immobilon-P PVDF membrane (Millipore) and incubated overnight at 4C with primary antibody diluted in TBS (Fisher) supplemented with 0.1% Tween20 (Fisher) and either 5% BSA (Cell Signaling) or 5% non-fat dry milk (Oxoid). Blots were then incubated at room temperature for 1 hour with the relevant secondary antibody diluted in TBS supplemented with 0.1% Tween20 and 5% non-fat dry milk and then detected using ECL Western Blotting Substrate (Pierce). Protein transfer, detection, and quantification using LiCor was performed as previously described^9^.

### Tracing of Indels by Decomposition (TIDE) analysis

Genomic DNA was extracted from cell pellets using the DNeasy Blood & Tissue kit (Qiagen). An approximately 800-bp region centered on the sgRNA cut site was PCR amplified from 50 ng of genomic DNA using primer pair P5 (sgGFP) or P6 (sgRPA1-1) (**Supplementary Table S1**). Completed PCR reactions were treated with exonuclease I (NEB) according to the manufacturer’s protocol and then purified with the QIAquick PCR purification kit (Qiagen). 20 ng of purified PCR product was Sanger sequenced using an M13-forward primer and chromatograms were analyzed as previously described^34^.

### Flow cytometry analysis for ***in vitro* experiments**

A549-SpCTRE or A549^GFP^-SpCTRE cells for flow cytometry analysis were collected using TrypLE Express (Thermo) according to the manufacturer’s protocol to preserve CD4^T^ cell surface expression. Approximately 1 million cells were resuspended in PBS containing human TruStain FcX (Biolegend) and incubated at 4C for 10 minutes. Cells were stained with the α-CD4 antibody for 30 minutes at 4C, then washed twice with PBS and resuspended in PBS containing 1 µg/mL DAPI. All α-CD4 staining was performed with PE anti-human CD4 antibody clone RPA-T4 (Biolegend), unless otherwise stated. For analysis of GFP expression in CD4 negative cells (i.e. A549^GFP^-lentiCas9-Blast), approximately 1 million cells were washed twice with PBS and resuspended in PBS containing 1 µg/mL DAPI. All flow analysis was performed on an LSR Fortessa or LSR II (BD Biosciences).

### CD4^T^ extracellular domain analysis

A549 cells were co-transfected with a 50:50 mix of each CD4 domain variant and pLenti Neo CMV/TurboGFP using JetPrime transfection reagent (Polyplus) at a 2:1 JetPrime:DNA ratio. Media was changed 24 h after transfection and cells were selected with puromycin for 3 days. Flow cytometry analysis was performed as described above with PE anti-human CD4 antibody clones M-T466 (Miltenyi), RPA-T4, SK3, and OKT4 (Biolegend). Mean fluorescence intensity (MFI) of GFP positive cells was calculated using FlowJo. To compare antibody binding between CD4^T^ and full-length CD4, A549 was transduced with pLenti CMV Puro/CD4 lentivirus and selected with puromycin for 3 days. This cell lines was mixed with A549^GFP^-SpCTRE and flow cytometry analysis was performed as described above using anti-human CD4 antibodies RPA-T4 (APC) and OKT4 (PE).

### Magnetic cell separation (MACS)

A549-SpCTRE cells were spiked into an A549^GFP^ cell suspension at an abundance of 5-20%. This mixture was incubated with CD4 microbeads (Miltenyi) and subjected to magnetic separation with LS columns (Miltenyi) according to the manufacturer’s protocol. Eluted cells were cultured for 3 days to allow for dissociation of magnetic beads and then collected for flow cytometry analysis of CD4^T^ purity, as described above. Results represent three independent biological replicates.

### Transduction and enrichment of SpCTRE PDXs

Established PDX tumors were resected and dissociated to a single cell suspension using a gentleMACS tissue dissociator with a human tumor dissociation kit (Miltenyi). Red blood cells were lysed with ACK lysing buffer (Lonza) and mouse stroma cells were subsequently removed by negative magnetic bead selection using a mouse cell depletion kit (Miltenyi). Cells were then transduced *ex vivo* with pSpCTRE lentivirus at a relative MOI of 1-6 (based on a functional titer performed in A549) in the presence of 48 µg/mL polybrene in a swinging bucket rotor for 30 min at 800 x g. After transduction, cells were washed twice with PBS to remove the lentivirus and polybrene and engrafted in a 50% Matrigel (BD) mixture into a single flank of 6-8 week old NSG (Jax) or Nude (Envigo) mice. Resulting tumors were dissociated to a single cell suspension, red blood cells were lysed, and mouse stroma was removed as described above. Cells were prepared for flow cytometry sorting or analysis by resuspending 5-10 million cells (sorting) or 1 million cells (analysis)No in FACS buffer (PBS with 2% FBS, 1X pen/strep and 1 mM EDTA) containing human TruStain FcX (Biolegend) and mouse TruStain FcX (anti-mouse CD16/32, Biolegend) and incubating at 4C for 10 minutes. Cells were stained with APC/Cy7 anti-human CD4 (clone RPA-T4, Biolegend) and AlexaFluor647 anti-mouse H-2K^d^ (clone SF1-1.1, Biolegend) for 30 minutes at 4C. Cells were washed twice with MACS buffer (PBS with 0.5% BSA) and then resuspended in sorting buffer (PBS with 0.5% BSA, 2.5 mM MgCl_2_, 0.5 mM CaCl_2_, 1 µg/mL DAPI and 100 U/mL DNaseI, NEB) and incubated for 30 minutes at room temperature before sorting. CD4^T^ positive, H-2K^d^ negative, DAPI negative cells were collected using a FACSAria (BD Biosciences) and engrafted in a 50% Matrigel mixture into a single flank of a 6-8 week old NSG or Nude mouse. Resulting tumors were collected and analyzed for CD4^T^ expression as described above. If the tumor was less than 50% CD4^T^ positive, it was sorted again as described and if the tumor was >=50% CD4^T^ positive it was propagated or cryo preserved in RPMI supplemented with 10% FBS, 1x pen/strep, and 10% DMSO. SNP profiles of SpCTRE PDXs were compared to pre-sorted samples using qPCR genotyping of 8 informative SNPs to confirm identity.

### *In vivo* competition assays in SpCTRE PDXs

Established SpCTRE PDX tumors were dissociated to a single cell suspension, red blood cells were lysed, and mouse stroma was removed as described above. Cells were then independently transduced *ex vivo* with sgTrack lentivirus at a relative MOI of 1 as described above. Each sgTrack transduced SpCTRE PDX was engrafted in a 50% Matrigel mixture into a single flank of a 6-8 week old NSG or Nude mouse. Resulting tumors were dissociated to a single cell suspension, red blood cells were lysed, and mouse stroma was removed as described above. Cells containing control and test sgRNAs were mixed with an equal ratio of GFP and mCherry positive cells and engrafted in a 50% Matrigel mixture into a single flank of 6-8 week old NSG or Nude mice. Once tumors reached ∼100 mm^3^, mice were randomized to control or dox-treated groups, with dox-treated mice receiving 625 mg/kg doxycycline chow (Envigo). Tumors were collected once they reached 1000 mm^3^, dissociated to a single cell suspension, and prepared for flow cytometry analysis or sorting as described above. Indel analysis of sorted cells was performed as described above and fitness scores and log ratios were calculated as described in **Supplementary Fig. 4b**. The Wilcoxon rank sum statistic was used to test if the fitness scores in the dox-treated group were smaller than the fitness scores in the control group.

### Recombinant AAV production and validation

Recombinant AAV2/6 pseudotyped virus was produced by the Boston Children’s Hospital Viral Core. PC9 cells transduced with lentiCas9-Blast and selected with blasticidin for 7 days were subsequently transduced with rAAV at an MOI of 2.3. Control and rAAV transduced PC9-lentiCas9-Blast cells were treated with 1 µM osimertinib (Selleck Chemicals) or DMSO (Corning) until they reached confluency, at which time cell pellets were collected for genomic analysis, described below. Osimertinib resistance was confirmed in cells previously transduced with rAAV and selected with osimertinib by seeding 3000 viable cells per well in 100 µL/well media containing a dilution series of osimertinib. Viability was assayed 72 h after plating using CellTiter-Glo (Promega).

### *In vivo* homology-directed repair in MSK-LX29

Crizotinib (LC Laboratories), osimertinib (LC Laboratories), or the combination were formulated in 0.5% hydroxypropyl methylcellulose (HPMC) and given by oral gavage at 25 mg/kg daily for 5 days per week. An MSK-LX29-SpCTRE tumor from a mouse fed with dox chow for 4 weeks prior to dissociation was incubated with the rAAV at an MOI of 1.6 million genomic copies/cell for 1 hour at 37C. Cells were washed twice with PBS and then engrafted in a 50% Matrigel mixture in the flank of 6-8 week old NSG mice. All mice were administered dox chow at the time of engraftment and treatment groups were randomized once tumors reached 100 mm^3^. Tumors were collected once they reached 1000-1500 mm^3^ and genomic analysis was performed as described below.

### Genomic analysis of EGFR homology-directed repair

Genomic DNA was purified from dissociated tumors or cell pellets using the Gentra Puregene Cell kit (Qiagen). A 1,122 bp amplicon, which spans outside the rAAV homology arms to ensure amplification occurs from genomic DNA, was PCR amplified from 200 µg genomic DNA using primer pair P7. The PCR product was then digested with exonuclease I to remove excess primers, column purified using the QIAquick PCR purification kit (Qiagen), and 100 µg was used as the template for a second PCR with primer pair P8 to amplify a 273 bp region centered on the repair site. The second round PCR product was column purified and paired-end Next-Gen sequencing (NGS) was performed by the CCIB DNA Core Facility at Massachusetts General Hospital (Cambridge, MA). NGS sequencing was analyzed using the R statistical computing environment to determine the proportion of reads that underwent rAAV-mediated homology-directed repair.

## ACKNOWLEDGEMENTS

We thank the MSKCC Flow Cytometry core for their technical assistance, members of the Antitumor Assessment Core Facility for assistance with *in vivo* experiments, and all members of the Rudin lab for critical comments. We thank the Center for Computational and Integrative Biology (CCIB) at Massachusetts General Hospital for the use of the CCIB DNA Core Facility (Cambridge, MA) and the Boston Children’s Hospital Viral Core (Core grant 5P30EY012196, NEI). Supported by NIH U01 CA199215, U24 CA213274, P01 CA129243, and P30 CA008748 (CMR and JTP).

## AUTHORSHIP CONTRIBUTIONS

C.H.H. performed most of the experiments described here and drafted the manuscript. E.A.C. performed key in vitro assays including editing of GFP in A549^GFP^ cells and of EGFR in PC9 cells. E.dS. directs the xenograft core facility in which the EGFR gene editing *in vivo* under drug selection were performed. N.S.S. and A.Q.V. provided technical assistance. G.H. provided statistical analysis. C.M.R. and J.T.P. supervised the project and helped draft the manuscript. All authors contributed to editing the manuscript.

## DATA AVAILABILITY

No datasets were generated or analyzed during the current study.

---

**Supplementary Figure 1.**
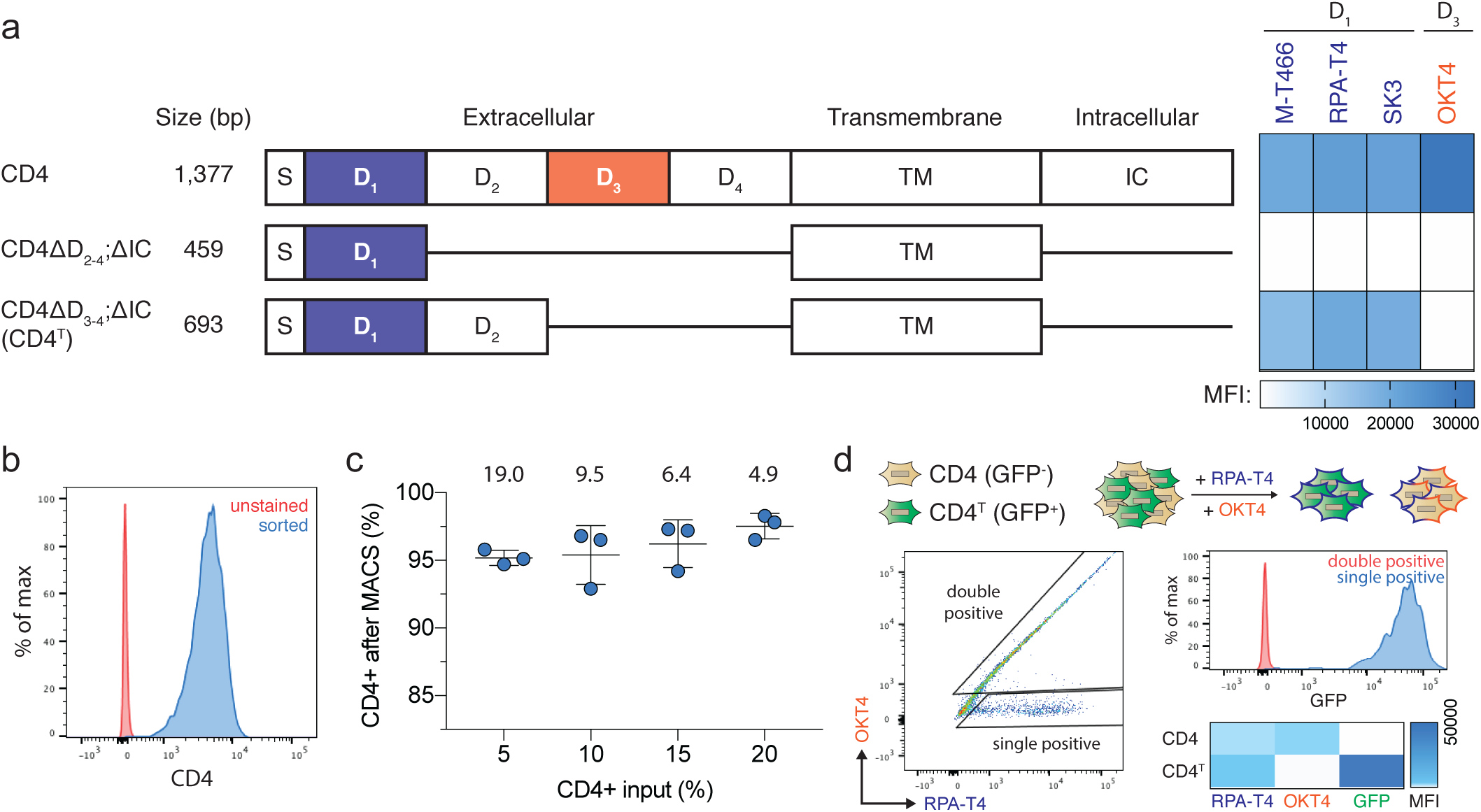
Truncated CD4^T^ is a size-efficient selectable marker for flow cytometry and magnetic bead selection. (**a**) Domain structure of wildtype human CD4 and truncated constructs CD4ΔD_2-4_;ΔIC and CD4ΔD_3-4_;ΔIC (hereafter CD4^T^), where the indicated deletions are replaced by flexible linkers. Heatmap depicts flow cytometry staining intensity of commercially available α-CD4 antibodies, which target the indicated extracellular domains of CD4, to the indicated CD4 constructs. S, signal peptide; TM, transmembrane domain; IC, intracellular domain; MFI, mean fluorescence intensity (**b**) Flow cytometry analysis of A549 cells expressing CD4^T^ and enriched by fluorescence activated cell sorting (FACS). (**c**) Magnetic bead enrichment of A549 cells expressing CD4^T^, with the indicated enrichment factors (above). Error bars are SD (n = 3). (**d**) α-CD4 staining strategy with domain D_1_ and D_3_ targeting antibodies to differentiate CD4^T^ and full-length CD4 using flow cytometry. A mixture of GFP-negative cells expressing full length CD4 and GFP-positive cells expressing CD4^T^ were stained with the indicated α-CD4 antibody clones. Double positive cells were exclusively GFP-negative (expressing full length CD4) and cells single positive for the domain D_1_ targeting α-CD4 antibody were exclusively GFP-positive (expressing CD4^T^).

**Supplementary Figure 2.**
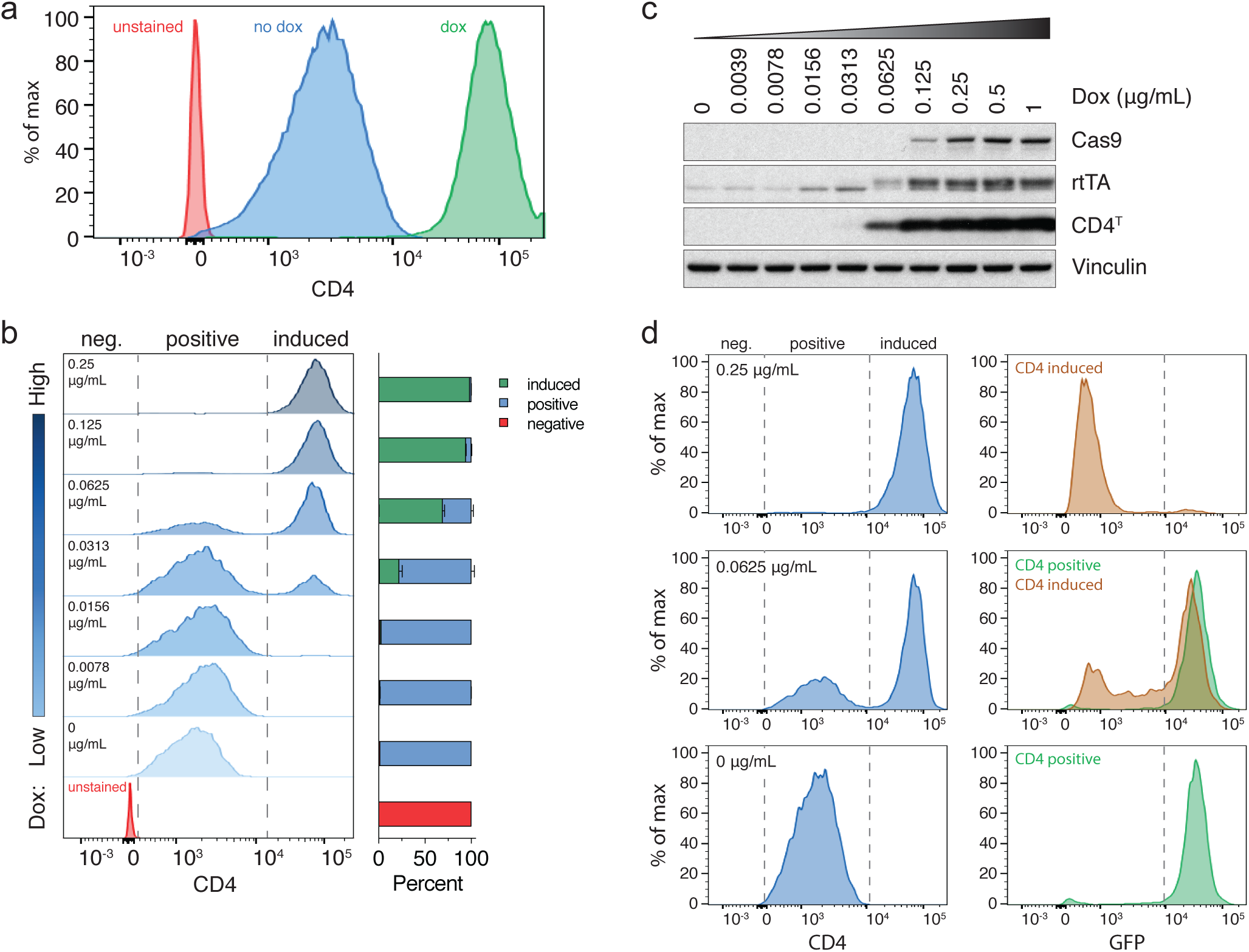
Dox induction of EFS promoter activity and CD4^T^ expression above basal levels is a marker for Cas9 expression and genome editing. (**a**) Flow cytometry analysis of CD4^T^ expression in A549-SpCTRE cells treated with (green) or without (blue) dox. (**b**) Representative flow cytometry analysis of CD4^T^ expression with the indicated dox concentrations in A549-SpCTRE cells. Percentage of cells in the negative, positive, and induced gates is displayed in the bar graph (right). Error bars are SD (n = 3), SD < plotting character not drawn. (**c**) Representative Western blot analysis of experiment in panel **b**. (**d**) GFP editing of A549^GFP^-SpCTRE cells with sgGFP based on gating of CD4^T^ positive or induced populations at the indicated dox doses. Loss of GFP expression is only detected in cells that induce CD4^T^. Data is representative of three independent experiments.

**Supplementary Figure 3.**
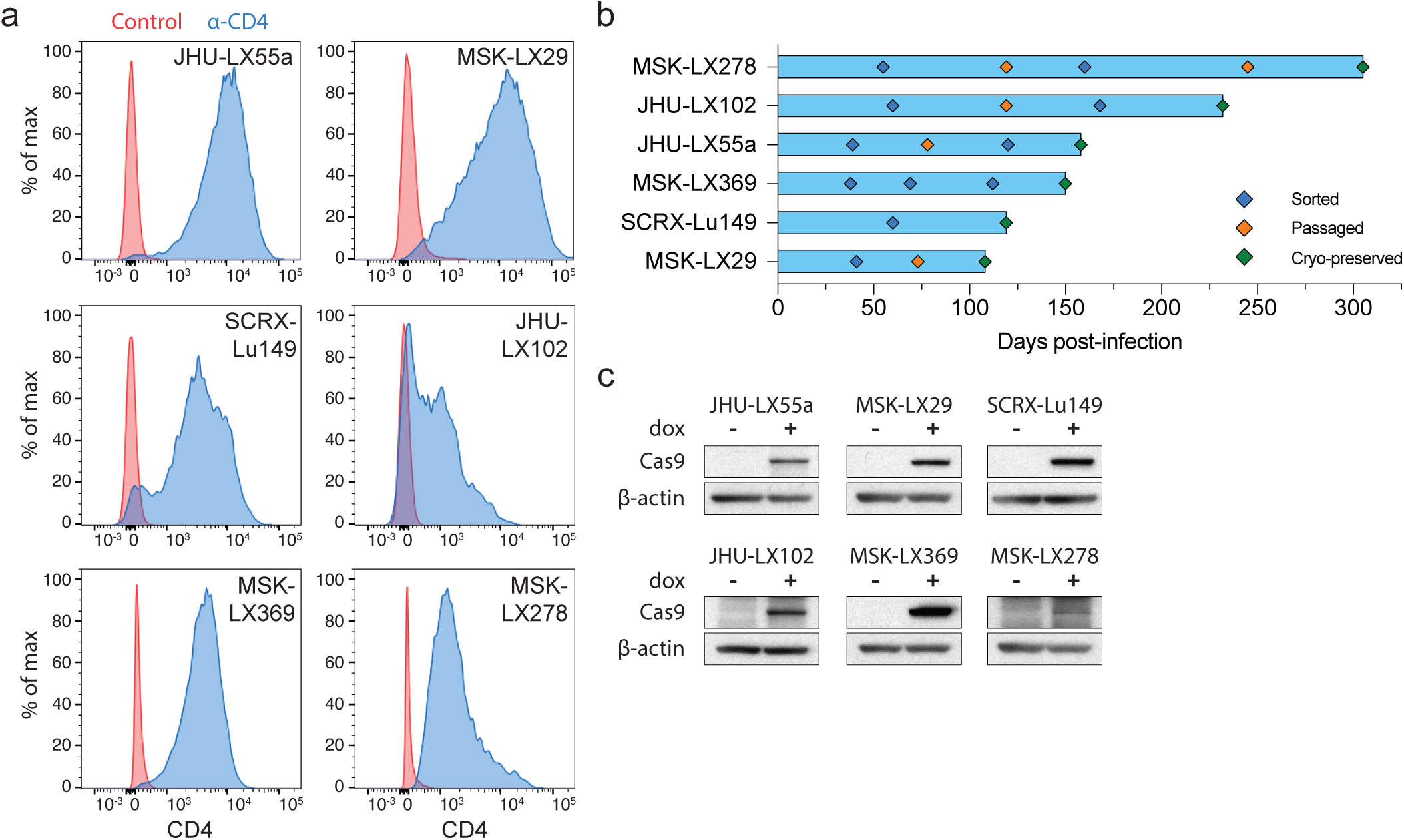
Generation of Cas9-expressing PDXs using pSpCTRE. (**a**) CD4^T^ staining of SpCTRE PDXs successfully transduced with pSpCTRE lentivirus and enriched for CD4^T^ positive cells using FACS. Control is fluorescence minus one (FMO) that excludes α-CD4 antibody. (**b**) Timeline for SpCTRE PDX model generation. PDXs were 1) sorted to enrich for CD4^T^ positive cells if CD4^T^ positive percentage was below 50% (blue), 2) passaged to expand for additional sorting or cryopreservation (orange), or 3) cryopreserved (green). (**c**) Western blot analysis of Cas9 expression in the indicated pSpCTRE PDXs from control or dox-treated mice.

**Supplementary Figure 4.**
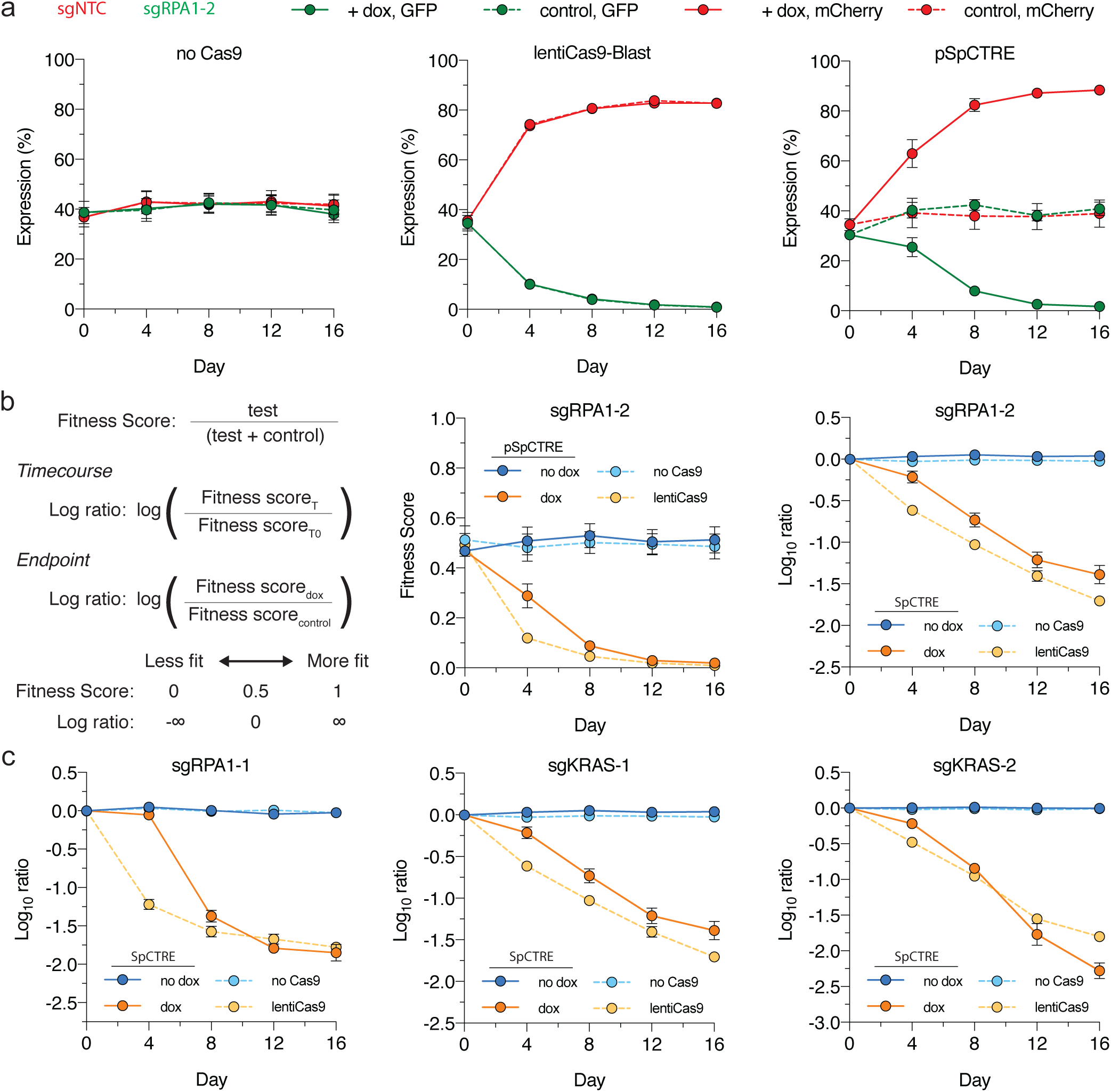
Competition assay utilizing sgTrack vectors effectively determines fitness effects of gene disruption. (**a**) Competition assay of sgRPA1-2 in A549 with no Cas9, constitutive Cas9 expression from lentiCas9-Blast, or dox-inducible Cas9 expression from pSpCTRE (left, middle and right panels, respectively). Competition assays were performed with (solid) or without (dashed) dox. Error bars are SD (n = 3), SD < plotting character not drawn. (**b**) Statistical analysis of the competition assay experiment in panel **a**. Fitness score calculates the percent of single positive fluorescent cells with the test sgRNA relative to the total number of single positive fluorescent cells representing both the test and control sgRNAs (coupled here with GFP or mCherry expression, respectively). The log ratio for a timecourse experiment then compares the fitness score at each timepoint to the initial fitness score at T0 or, for endpoint analysis, it compares the fitness score between control and dox-treated groups. A negative log ratio indicates the test sgRNA reduced the fitness of the cells relative to the control sgRNA. Solid lines represent competition assays in A549-SpCTRE cells either with (orange) or without (blue) dox. Dashed lines represent competition assays in A549 with (orange) or without (blue) Cas9 expression. Error bars are SD (n = 3), SD < plotting character not drawn. (**c**) *In vitro* competition assays in A549 with sgRPA1-1, sgKRAS-1, and sgKRAS-2 (left, middle and right panels, respectively). Log ratio calculations and line assignments are as described in panel **b**. Error bars are SD (n = 3), SD < plotting character not drawn.

**Supplementary Figure 5.**
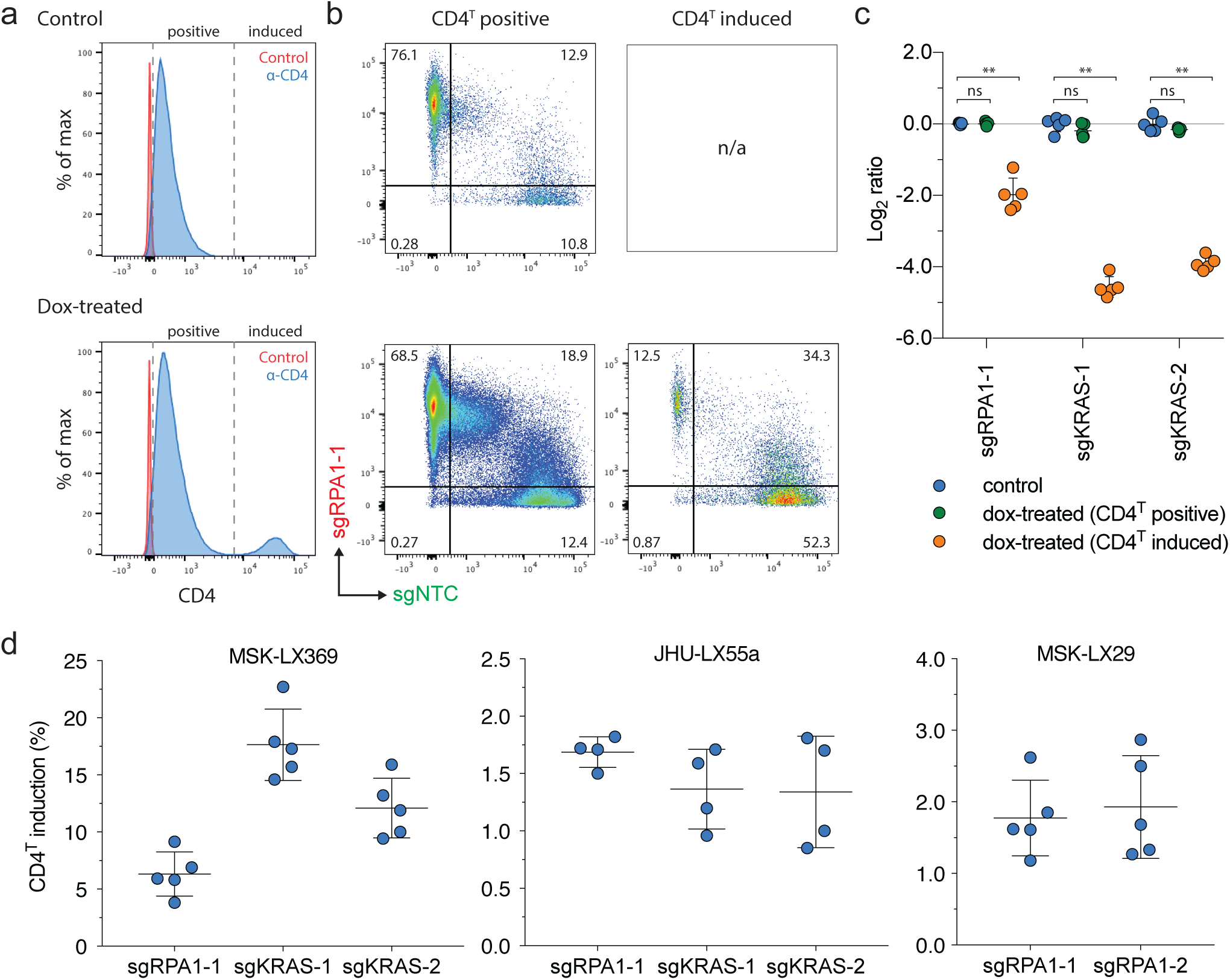
CD4^T^ induction in SpCTRE PDXs and gating strategy for *in vivo* competition assay analysis. (**a**) Representative flow cytometry analysis of CD4^T^ induction for the competition assay with sgRPA1-1 in MSK-LX369. (**b**) Flow cytometry analysis of sgRPA1-1 (mCherry) and sgNTC (GFP) in tumors from panel **a**. Cells are first gated on CD4^T^ positive or CD4^T^ induced populations as shown in panel **a**. (**c**) Log ratio of MSK-LX369 sgRPA1-1 competition assay for control or dox-treated mice gated on either CD4 positive or CD4 induced populations. Error bars are SD (n = 5), SD < plotting character not drawn. Each data point represents one tumor. ns = p > 0.05 and ** p < 0.01 between groups by Wilcoxon rank sum test. (**d**) Percent of CD4^T^ induced cells from dox-treated mice for competition assays with the indicated sgRNAs in the SpCTRE PDXs MSK-LX369, JHU-LX55a, and MSK-LX29 (left, middle, and right panels, respectively). Error bars are SD. Each data point represents one tumor.

**Supplementary Figure 6.**
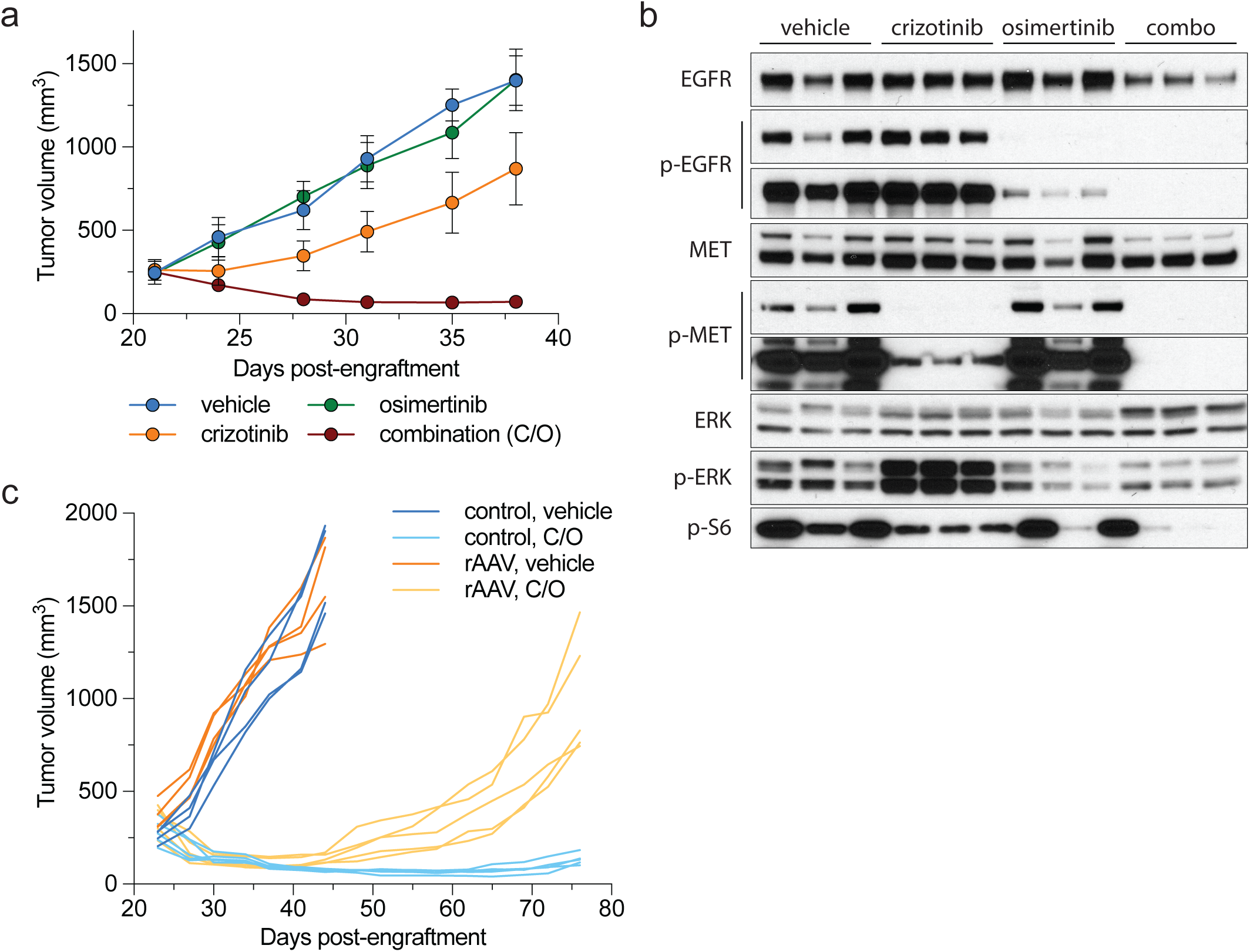
Evaluation of EGFR inhibitor combination therapy in a *MET* amplified PDX (**a**) MSK-LX29 tumor growth curves for the indicated treatment arms. Error bars are SD (n = 3), SD < plotting character not drawn. C/O, crizotinib/osimertinib combination. (**b**) Western blot analysis of the tumors treated in panel **a**. (**c**) Spider plot of the tumor growth experiment shown in **Fig. 3b**.

**Supplementary Figure 7.**
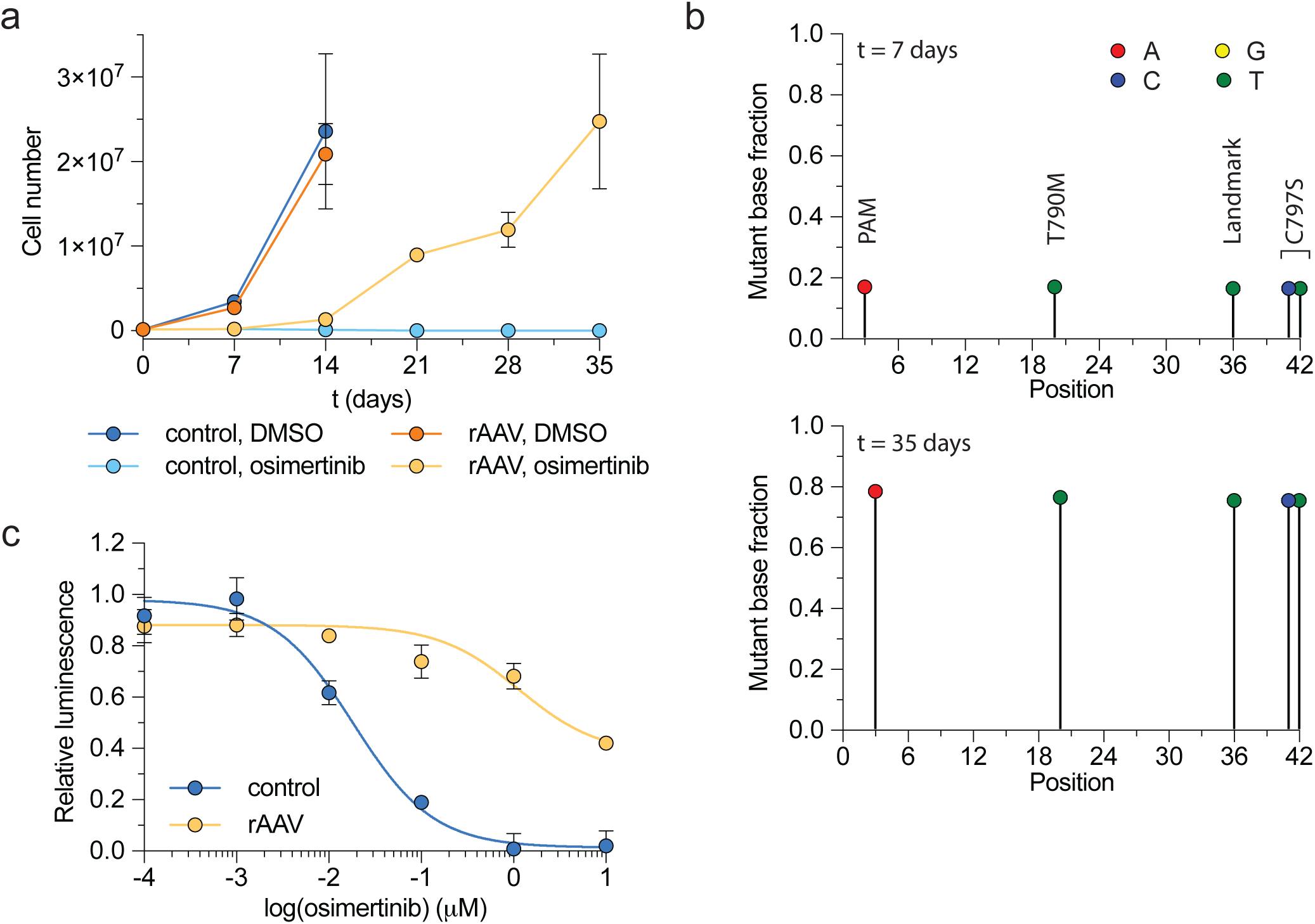
Evaluation of acquired osimertinib resistance driven by rAAV-mediated homology-directed repair in PC9 cells *in vitro*. (**a**) Cell line growth curves of PC9 cells under the indicated conditions. Error bars are SD (n = 3), SD < plotting character not drawn. (**b**) Sequencing analysis of PC9 cells from the rAAV, osimertinib group at t = 7 and 35 days. (**c**) Osimertinib dose response curves of PC9 cells from the rAAV, osimertinib or control, DMSO treatment arms from panel **a**. Error bars are SD (n = 3), SD < plotting character not drawn.

